# APRIL and BAFF increase breast cancer cell stemness

**DOI:** 10.1101/151902

**Authors:** Vasiliki Pelekanou, George Notas, Paraskevi Athanasouli, Konstantinos Alexakis, Fotini Kiagiadaki, Nikolaos Peroulis, Konstantina Kalyvianaki, Errika Kampouri, Hara Polioudaki, Panayiotis Theodoropoulos, Andreas Tsapis, Elias Castanas, Marilena Kampa

**Affiliations:** Laboratory of Experimental Endocrinology, School of Medicine, University of Crete, Heraklion, GR-71003, Greece; Present affiliation: Department of Pathology, School of Medicine, Yale University, New Haven, CT 06510, USA; Department of Biochemistry, School of Medicine, University of Crete

**Keywords:** APRIL, BAFF, TNF members, pluripotency, EMT, cancer stem cells, therapy resistance, aromatase inhibitors

## Abstract

Recent advances in cancer immunology revealed immune-related properties of cancer cells as novel promising therapeutic targets. The two TNF superfamily members, APRIL and BAFF even though were primarily studied in lymphocyte maturation, they have also been associated with tumor growth and aggressiveness in a number of solid tumors, including breast cancer. In the present work we studied the effect of APRIL and BAFF on epithelial to mesenchymal transition and migration of breast cancer cells, and their action on the sub-population of cancer stem cells identified by autofluorescence and ALDH activity. Their action on an number of pluripotency genes was examined and breast cancer stem cell ability to form mammospheres was also utilized. The receptor and the signaling pathway involved as well as the role of steroid hormones in their action were also investigated. Our findings show that both APRIL and BAFF increase epithelial to mesenchymal transition and migratory capacity of breast cancer cells, as well as cancer stem cell numbers, by inducing pluripotency genes such as KLF4 and NANOG. These effects are mediated by their common receptor BCMA and the JNK signaling pathway. Interestingly, androgens enhance APRIL transcription and subsequently its pluripotency effect. In conclusion, our data support the significant role of APRIL and BAFF in breast cancer disease progression and provide evidence for a new possible mechanism of therapy resistance, that could be particularly relevant in aromatase inhibitors-treated patients, were local androgen is increased.

## Introduction

The breast cancer 5-year survival has significantly increased, edue to early detection, and the introduction of therapies, such as specific estrogen receptor modulators, aromatase inhibitors, specific chemotherapeutic agents and more comprehensive recent precision medicine strategies. However, long (10-years) survival decreases dramatically to 78% (http://www.cancerresearchuk.org/health-professional/cancer-statistics/statistics-by-cancer-type/breast-cancer/survival#heading-Zero), suggesting that, in spite of our progress in breast cancer cell biology, other elements are currently uncovered or understood.

Cancer stem cells are key players in tumor development, progression and metastasis as well as resistance to therapy. This cell population displays potential to proliferate infinetely, to initiate and/or reform a tumor and an enhanced resistance to therapy ^1^. Adult stem cells have been identified in the normal mammary gland development and function see ^2, for a recent review^, but also exert specific dynamics in in various mammary pathologies ^3^. Breast stem cells interact in their niche with surrounding stromal and epithelial cells, through specific molecular crosstalk pathways ^4,5^. These cancer-immune cell interactions are mostly orchestrated by tumor microenvironment ^6,7^. However, established immune-related therapies target immune cells (resident or infiltrating the tumor stroma) ^8^, leading to an immune checkpoint blockade ^9^, while the notion of immune-related properties of the cancer cell *per se* and its possible regulation as a possible therapeutic target is less well defined ^10,11^.

A number of immune-related molecules are involved in the above mentioned interactions and being targeted in tumor immunotherapy approcahes. Among them, TNF superfamily members (including TNF, FAS and TRAIL and their receptors) ^8^, have been actively investigated and targeted in a number of malignancies. The TNF superfamily includes 19 different ligands and 29 receptors, controls cell survival and differentiation and plays an important role in the growth, organization, and homeostasis of different tissues, by modulating major signaling pathways ^12^. Our group has focused on two members of this superfamily, namely APRIL (A PRoliferation Inducing Ligand) and BAFF (B-cell Activating Factor of the TNF Family). These two ligands, act via two common receptors, B-Cell Maturation Antigen (BCMA) and Transmembrane Activator and CAML Interactor (TACI), while additionally, BAFF-Receptor (BAFF-R) is a specific receptor for BAFF. They had initially been reported to have a central role in lymphocyte maturation; however, they have been also identified as significant players in several other conditions including neoplasia ^13^. BAFF and APRIL have been detected in solid tumors ^14^, activating significant kinase signaling pathways, such as p38, JNK, NFκB and inducing, in the majority of cases, cell survival and growth. Previously, we have shown BAFF and/or APRIL presence in many normal and cancer solid tumors, including breast cancer ^15–17^, in which BAFF is constantly expressed in tumors, while APRIL is related to tumor grade ^17^. Recently, higher APRIL expression was shown in human triple negative carcinomas and APRIL was reported to induce cell proliferation both *in vitro* and *in vivo*, suggesting an association of APRIL signaling pathways with tumor aggressiveness ^18^. In the present work, we investigated the possible relationship of APRIL and BAFF with steroids, the important players in breast cancer, and the possible role of APRIL and BAFF in inducing breast cancer stemness.

## Results

### BAFF and APRIL expression in breast cancer cell lines - Regulation by extranuclear-acting androgen

We have assayed the presence of these TNFSF members in an ERα-positive (T47D) and a triple negative breast cancer cell line (MDA-MB-231 cells, named thereafter MDA cells). BAFF, APRIL, BCMA, and BAFFR mRNA was present in both cell lines, while TACI transcript was constantly absent (Figure 1A).

**Figure 1.**
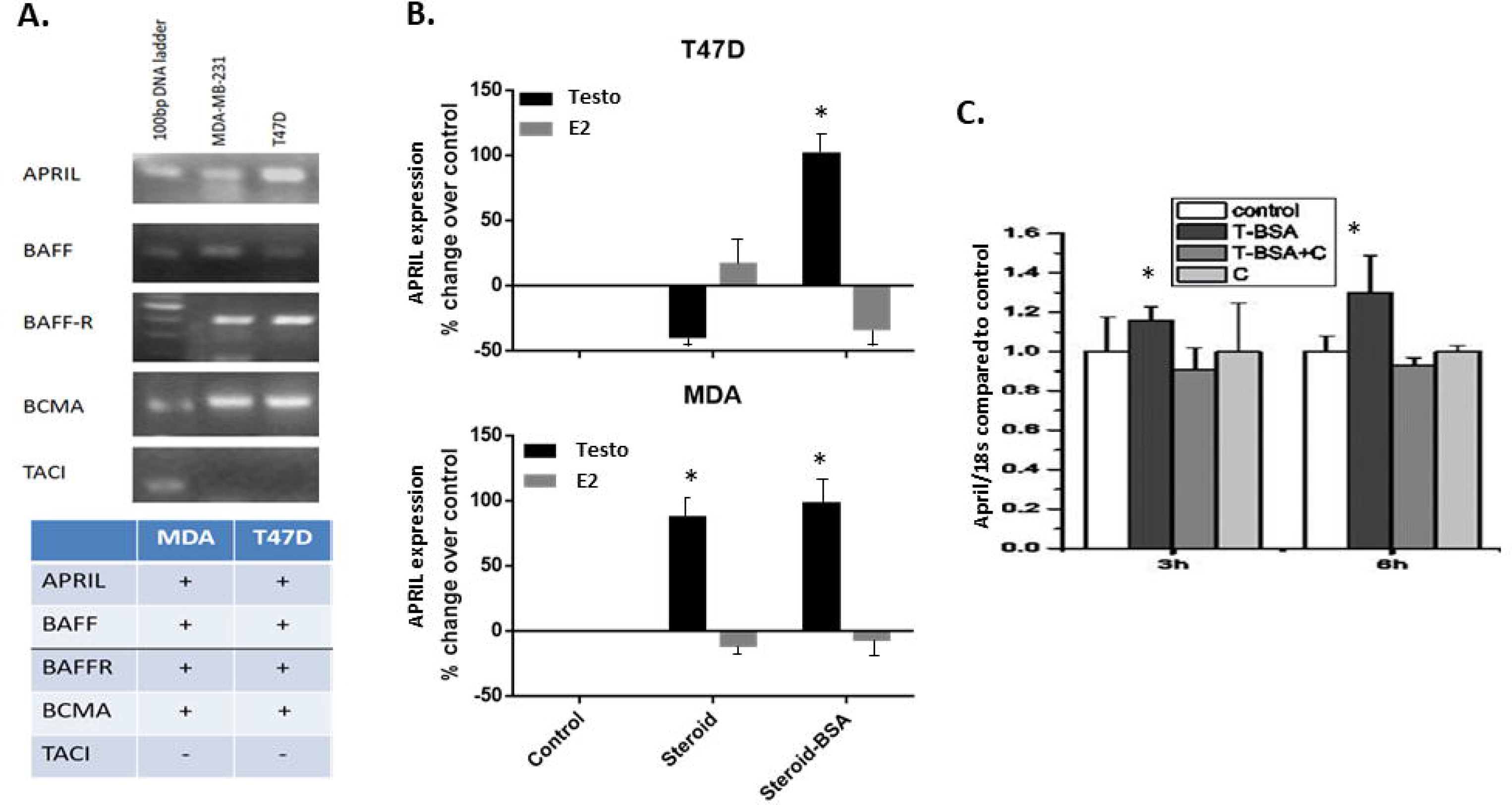
Expression levels of BAFF, APRIL, BAFF-R, BCMA and TACI in breast cancer cell lines and the ffect of steroid hormones on APRIL expression levels. **A.** The expression of BAFF, APRIL, BAFF-R, BCMA and TACI investigated by PCR in T47D and MDA-MB-231 cells breast cancer cell lines. **B.** APRIL expression (from Gene Chip analysis) after treatment of breast cancer cells for 3hrs with steroid hormones, testosterone (testo) or estradiol (E2), unconjugated or in their membrane impermeable form conjugated to BSA (steroid-BSA). **C.** The effect of testosterone-BSA on APRIL expression (investigated by PCR) in T47D breast cancer cells in the presence or absence of cyproterone acetate (*denotes p<0.05).

To explore whether there is a relation between steroids and APRIL and BAFF production we have, at first interrogated our previously reported transcriptome data on the effect of androgen (GSE18146) and estrogen (GSE32666 and GSE32668) on T47D and MDA cells. As depicted in Figure 1B a significant increase of APRIL transcription was observed in both T47D and MDA cells after 3h-incubation with testosterone. Interestingly, this increase was observed with the membrane-only acting androgen (Testosterone-BSA) suggesting an extranuclear androgen action. In contrast, estrogen had a minimal effect on APRIL transcription in both cell lines. Conversely, no significant modification of APRIL-BAFF receptors was found (not shown).

Exploring the promoter region of APRIL, with the rVista V 2.0 web resource (https://rvista.dcode.org/), we have identified four putative androgen response elements in the promoter of the APRIL gene (Supplementary Figure 1), an element suggesting the implication of the classical activated AR. In view of this and the transcriptional activation of APRIL and BAFF by membrane-acting steroids, we explored the possible modification of APRIL transcription after cell incubation with testosterone-BSA. Incubation of T47D cells with testosterone-BSA resulted in a significant increase of APRIL gene product at 3 and 6 hours (Figure 1C). Interestingly, this effect was inhibited by the antiandrogen cyproterone acetate (which was ineffective *per se*).

### BAFF and APRIL increase cell migration, epithelial-mesenchymal transition (EMT) and stemness in epithelial breast cancer cells

APRIL has been reported to be associated with increased breast tumor growth and metastasis ^18^. In our cell lines both APRIL and BAFF enhance cell migration (the effect is more prominent after a 48h-incubation, Figure 2A). Additionally, the EMT cell status of breast cancer cells was enhanced after treatment with either APRIL or BAFF (100ng/ml) (Figure 2 B and C). This was accompanied by actin cytoskeleton rearrangements, characteristic of migrating cells (Supplementary Figure 2). It is to note that APRIL and BAFF induce similar modification of EMT markers and migratory activity, suggesting an interaction occurring through the same receptor or receptors.

**Figure 2.**
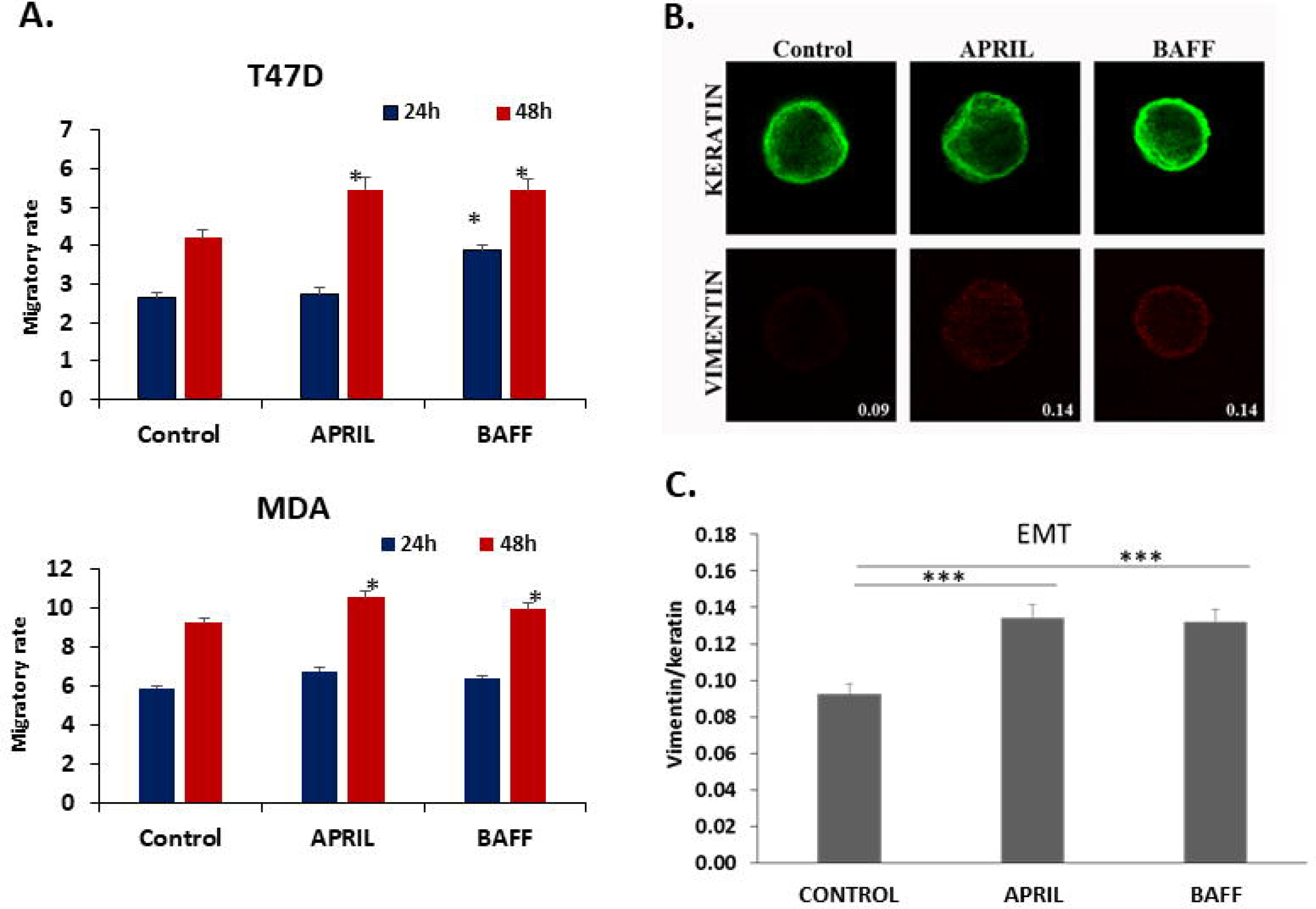
The effect of APRIL and BAFF on migration and EMT status. **A.** Effect of APRIL and BAFF (100ng/ml) on migration of T47D and MDA cells after 24 and 48 hours of treatment (expressed as percentage of control=100). **B.** Representative photos of control (untreated) T47D cells and APRIL or BAFF (100ng/ml) treated cells for 4 days immunostained for keratins (upper panel, green) and vimentin (lower panel, red). Fluorescence intensity was calculated using Image J. Values represent the ratio of vimentin/keratins expression in these cells. **C.** Graphical presentation of EMT status changes obtained by calculating the ratio of vimentin/keratins expression of 50 cells for each treatment condition. All samples were analyzed with t-test (***denotes p< 0.0001).

In addition to EMT, another factor necessary for metastatic potential and disease progression is a combination of genetic alterations and epigenetic events that recapitulate normal developmental processes including stem cell self-renewal ^19^ and the acquisition of “stem cell” properties, through a “dedifferentiation” process. T47D and MDA cells, treated with APRIL or BAFF (100ng/ml) exhibited an increased ability to form mammospheres. More specifically, the number of primary mammospheres increased by 40% after 9 days of incubation (Figure 3A). The effect of these two agents is more prominent in ERα-positive T47D than in triple negative MDA cells. Interestingly, testosterone-BSA per se increases mammosphere formation in T47D cells. When cells were incubated with testosterone-BSA and APRIL, an additive effect was observed (Figure 3B).

**Figure 3.**
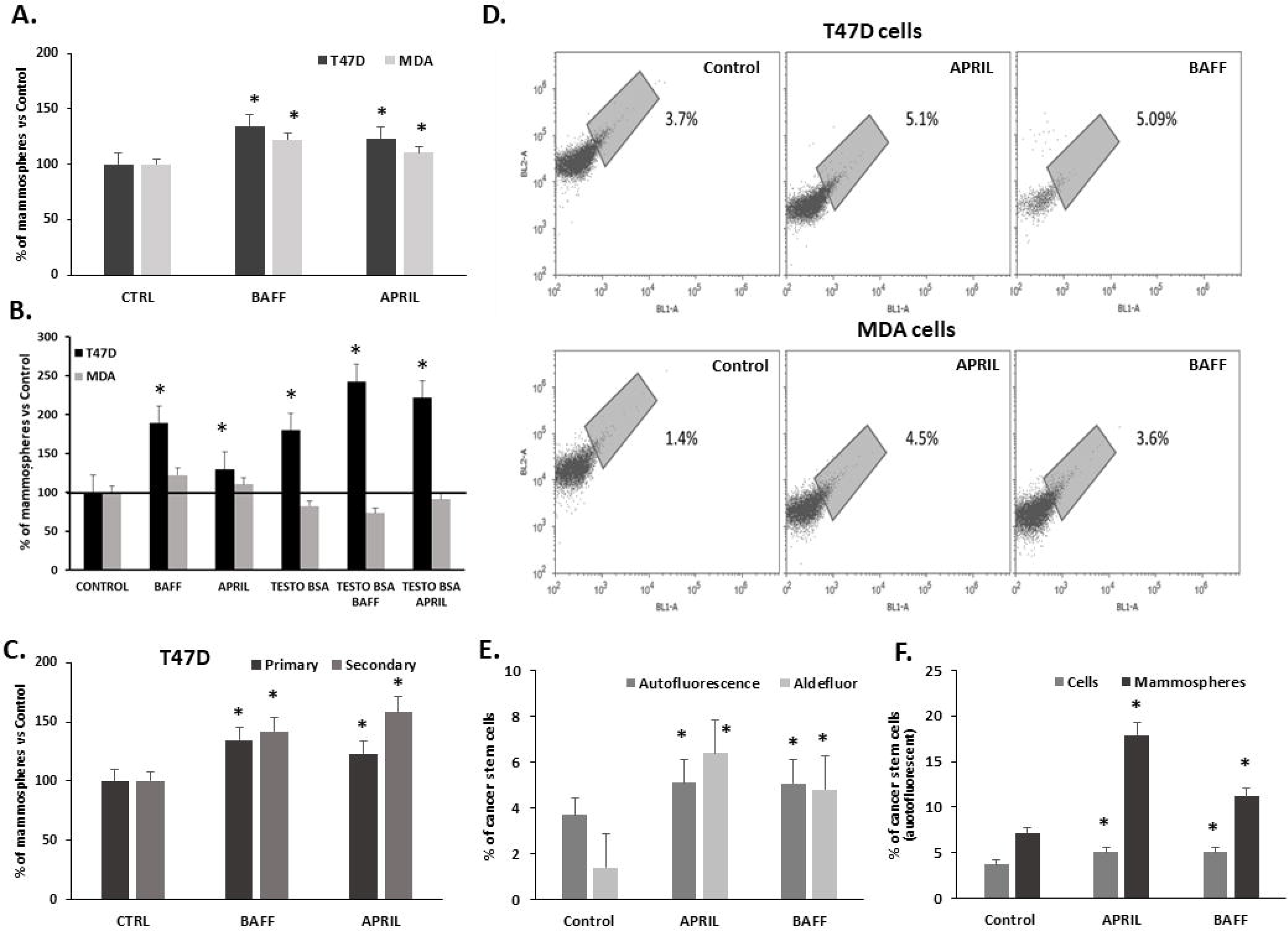
Effect of APRIL and BAFF on mammosphere formation and breast cancer stem cell population and testosterone on the effect of APRIL and BAFF on mammosphere formation. **A.** Percentage of mammospheres (primary) formed after treatment of breast cancer cells (T47D and MDA) with either BAFF or APRIL (100ng/ml) for 9 days. Results of three independent experiments are expressed as a percentage of control (untreated) cells (*denotes p<0.05). **B.** Percentage of mammospheres (primary) formed after treatment of breast cancer cells (T47D and MDA) with either BAFF or APRIL (100ng/ml) in the presence or absence of testosterone-BSA, for 9 days. Results of three independent experiments are expressed as a percentage of control (untreated) cells performed (*denotes p<0.05). **C.** Percentage of mammospheres formed after treatment of T47D cells with either BAFF or APRIL (100ng/ml) for 9 days (primary mammospheres) compared to secondary mammospheres and a subsequent 7-day period. Results of three independent experiments are expressed as a percentage of control (untreated) cells. (* P<0.05) **D.** Percentage of breast cancer stem cells (T47D and MDA) with or without treatment with BAFF or APRIL (100ng/ml) for 4 days as estimated by assaying them with high green autofluoresence in a BL2-A/BL1-A dot plot. Three independent experiments were performed and the results of a representative one is presented. **E.** Comparison of the percentages of breast cancer stem cells as estimated by autofluorescence in a BL2-A/BL1-A dot plot and ALDEFLUOR kit with or without treatment with BAFF or APRIL (100ng/ml) for 4 days. Three independent experiments were performed (*denotes p<0.05). **F.** Comparison of the percentages of breast cancer stem cells (identified by autofluorescence in a BL2-A/BL1-A dot plot) in T47D cells and mammospheres treated or not with BAFF or APRIL (100ng/ml) for 4 and 12 days respectively. Three independent experiments were performed (*denotes p<0.05).

We have particularly focused in the T47D cell line, in which mammosphere formation was more prominent. After dispersion of primary mammospheres and culture of cells under the same conditions (formation of secondary mammospheres) the effect of both cytokines was further enhanced, with APRIL being slightly more potent than BAFF (Figure 3C). The effect of BAFF and APRIL on the induction of stemness was further verified by the intrinsic autofluorescence of epithelial cancer stem cells, due to riboflavin accumulation in membrane-bounded cytoplasmic structures bearing ATP-dependent ABCG2 transporters ^20^. Incubation of breast cancer cells with APRIL or BAFF (100 ng/ml) for at least four days induced a significant increase in autofluorescence, from ~2% in control (untreated) cells to ~5% in APRIL or BAFF-treated cells Figure 3D). This increase in autofluorescence correlates and is of similar amplitude with the increase of ALDH1A1 activity, assayed by flow cytometry and the ALDEFLUOR^®^ kit (see Supplementary Figure 3 for an example and Figure 3E). Dispersed T47D cell mammospheres were assayed for a stem cell signature and they were found to have significantly more stem cells compared to cells from conventional monolayer culture (7.2% vs 2.1%). BAFF or APRIL treatment further increased this autofluorescent stem cell signature (17.9 vs 4.5 for APRIL and 11.2 vs 3.6 for BAFF respectively) suggesting that these cytokines have the capacity to induce pluripotency in T47D cells (Figure 3F).

To further verify the emergence of stemness after 4 days incubation with APRIL or BAFF (100 ng/ml), the transcription of pluripotency-related factors (SOX2, c-MYC, KLF4, NANOG and ALDH1A1) was investigated in T47D cells. The expression of the above genes was examined both at the mRNA (by Real Time PCR, Figure 4A) and protein level (by immunocytochemistry, Figure 4B). As shown, APRIL and BAFF mainly increase the expression of ALDH1A1, KLF4, and NANOG.

**Figure 4.**
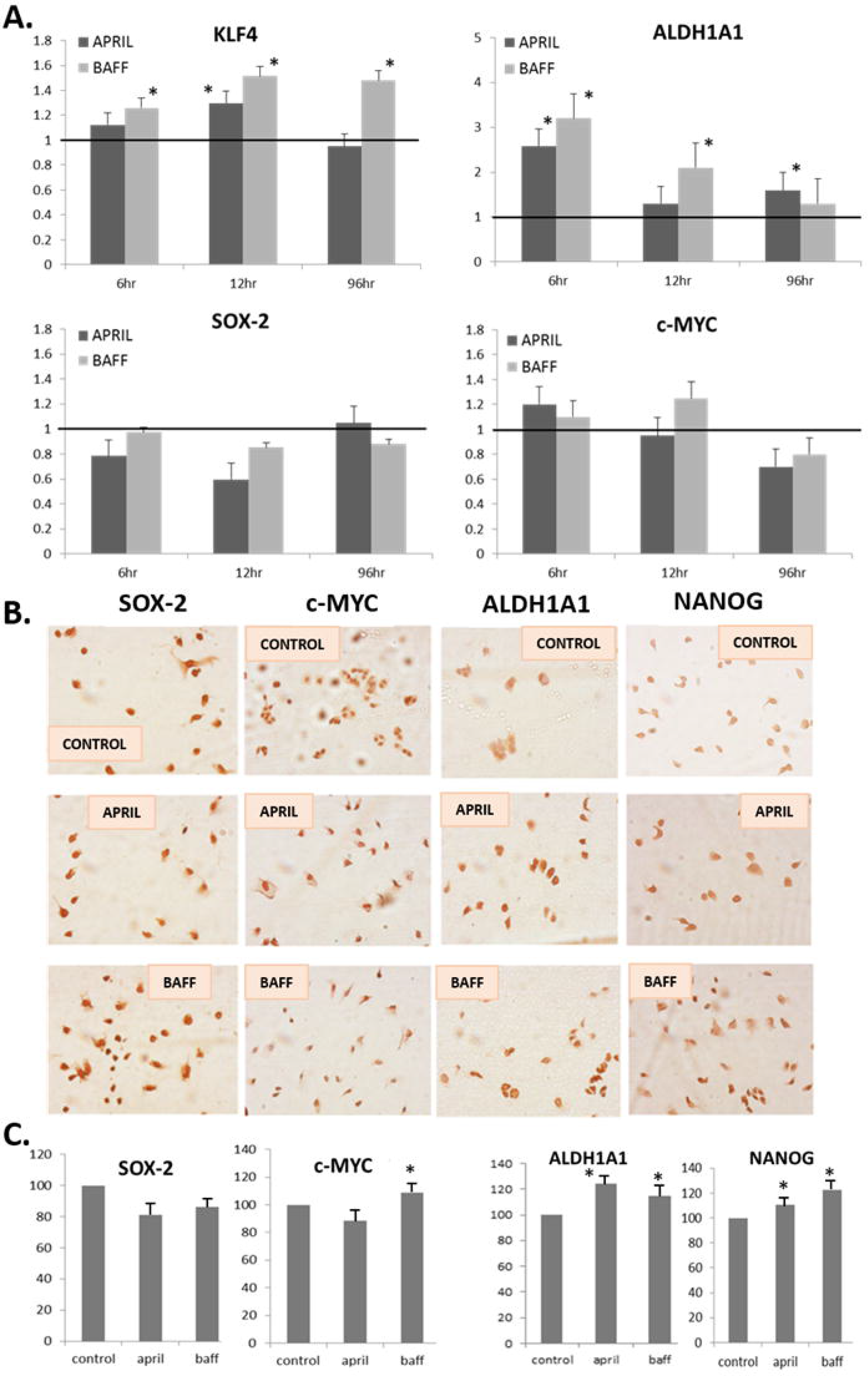
Expression levels of pluripotency markers in T47D cells after APRIL or BAFF treatment. **A.** Expression levels of ALDH1A1, KLF4, SOX-2, and c-MYC, genes (quantified by Real-Time PCR and expressed as percentage of control) after treatment of T47D cells with APRIL or BAFF (100ng/ml) for 4 days. Three independent experiments were performed (*denotes p<0.05). **B. and C.** Expression of SOX-2, c-MYC, ALDH1A1 and NANOG proteins (assayed by immunocytochemistry). Representative images of three independent experiments are presented (B) Quantification of the proteins’ expression levels was performed in at least 10 cells in three different photos for both control and treated cells using Image J and is expressed as percentage of control (C).

### APRIL and BAFF modulate stemness markers expression through BCMA and JNK signaling

Results presented so far suggest that APRIL and BAFF promote in a similar way the emergence of pluripotency in breast cancer cells. Taking into consideration that T47D cells express BCMA, on which both cytokines can bind, and BAFFR, an exclusive BAFF receptor (see Figure 1) ^12^, we have assumed that this effect might be mediated by BCMA. To confirm this hypothesis, we transfected T47D cells with a specific anti-BCMA siRNA. Transfected cells were then incubated with BAFF or APRIL and pluripotent/stem cells were assayed by their autofluorescence. As depicted in Figure 5A, cancer stem (autofluorescent) cells were significantly decreased, suggesting the mediation of this effect through BCMA. Here, we have explored the two pathways mediating post-receptor effects of BAFF and APRIL [TRAF signaling and NFκB activation (31), and BCMA-specific signaling, mediated by JNK (21)]: As shown in Figure 5B, T47D cells, transfected with a specific plasmid, carrying NFκB response elements in front of the firefly luciferase gene, do not show any activation after a 24h BAFF or APRIL incubation; contrariwise, the JNK specific inhibitor, SP600125 or shRNAs against JNK1 and JNK2, significantly blocked the effects of APRIL and BAFF on ALDH1A1 and KLF4 (Figure 5C and D), indicating that, in breast cancer cells, APRIL and BAFF binding to BCMA signal towards pluripotency is mediated by JNK1.

**Figure 5.**
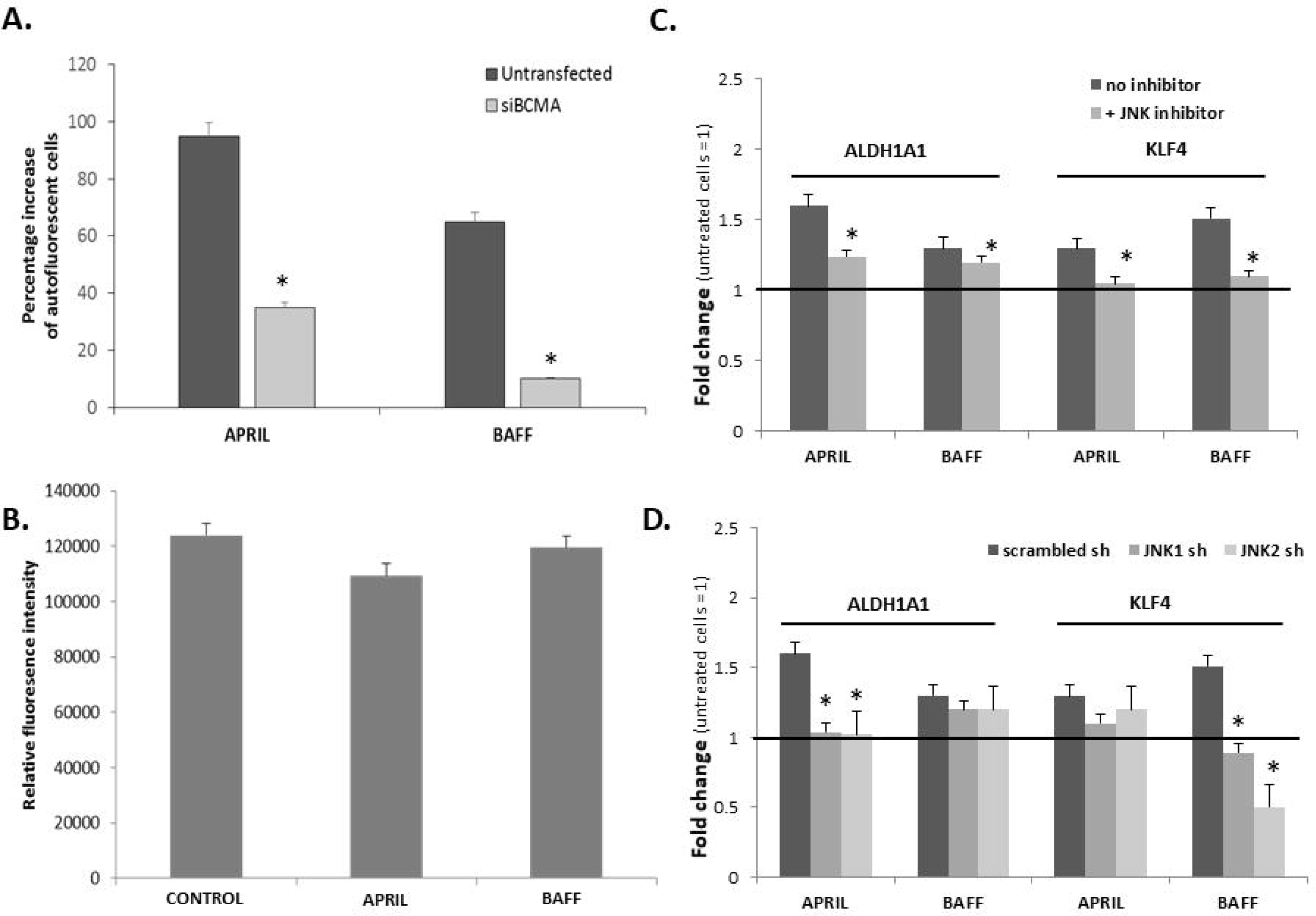
Effect of APRIL and BAFF on breast cancer stem cells in the presence of siBCMA. Percentage increase in the number of breast cancer stem cells transfected or untransfected with siBCMA and treated with BAFF or APRIL (100ng/ml) for 4 days. (Breast cancer stem cell number was estimated by counting the cells with high green autofluoresence in a BL2-A/BL1-A dot plot). **B. NFk-B activity** of untreated (control) and treated with APRIL or BAFF (100ng/ml, for 24hrs) T47D cells. Experiments were performed in triplicate. **C. and D. Effect of APRIL and BAFF on ALDH1A1 and KLF4 expression in the presence of a JNK inhibitor or sh RNA for JNK1 or JNK2.** Expression levels of ALDH1A1 and KLF4 after treatment of T47D cells with APRIL or BAFF (100ng/ml) for 4 days in the presence of a JNK inhibitor SP600125 (10μM) (C) or sh RNA for JNK1 or JNK2(D). Data are presented as a mean ±SEM of three independent experiments (*denotes p<0.05).

### APRIL in aromatase-inhibitor treated patients’ samples

As shown above, androgen enhances APRIL production in breast cancer cells regardless of estrogen receptor status. We hypothesized that increased local androgen (as could occur in patients treated with aromatase inhibitors that inhibit conversion of androgen to estrogen), could enhance APRIL production ultimately increase cancer pluripotency. We extracted APRIL data from the GEO-deposited GDS3116 study ^21,22^. In this series, transcriptome data of a series of 53 letrozole (an aromatase inhibitor) treated patients are included, prior and 3-months following therapy. 37/53 (69.8%) patients responded to the therapy (as estimated by changes in estrogen responsive genes and by ultrasound detected changes in tumor volume) while 16/53 (30.1%) were nonresponders. In responders, while in non-responders no change is observed (Figure 6A and B).

**Figure 6.**
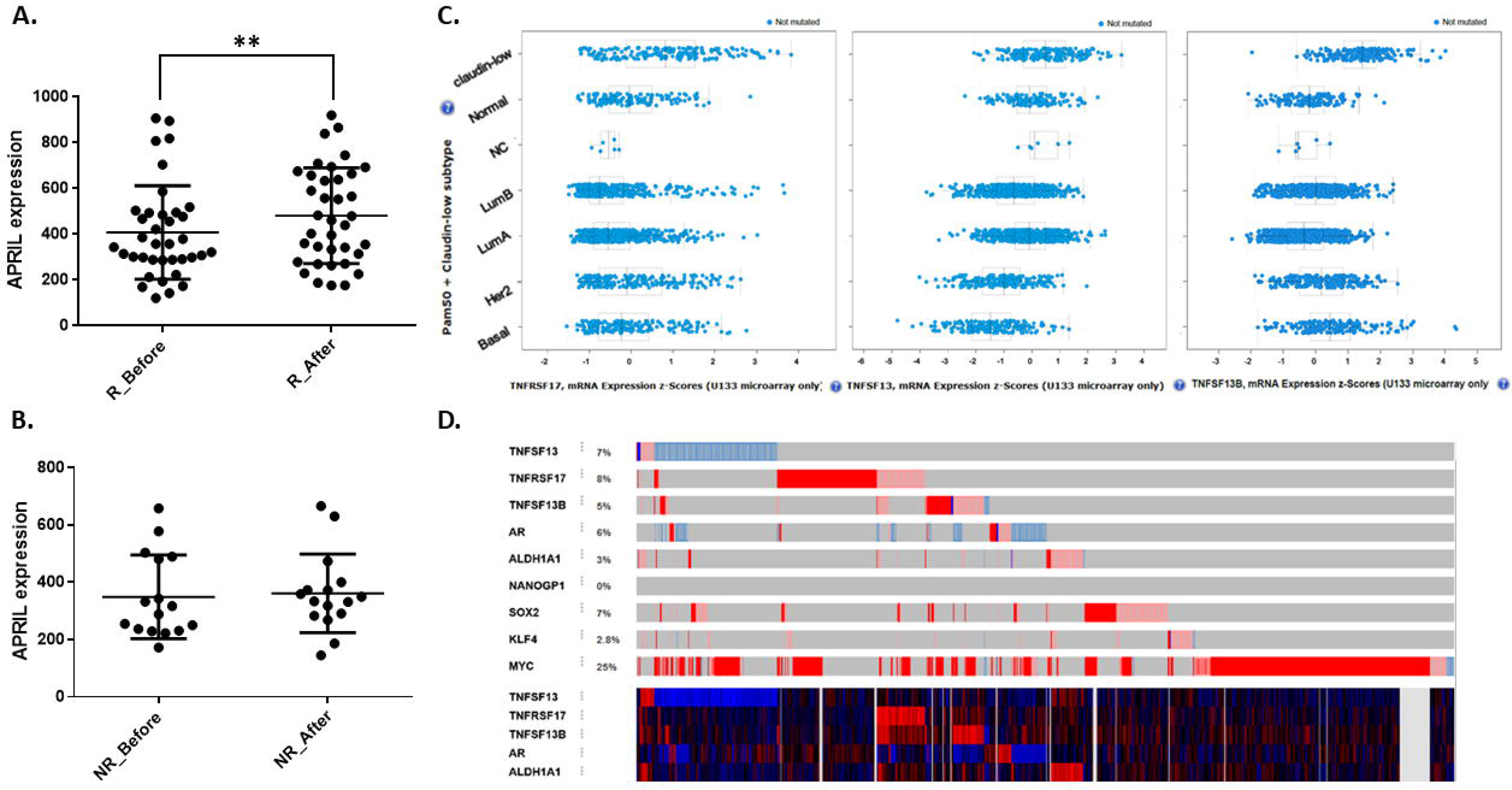
APRIL expression in breast cancer specimens from patients and Results of the analysis of the MetaBRIC study. IN cBioPortal^®^.**A.** Patients that have responded to letrozol therapy (R) and **B.** patients that have not responded (NR). The levels of APRIL expression are given before and after therapy. (**denotes p<0.001). **C.** The distribution of BCMA (TNFRSF17, left panel), APRIL (TNFSF13, middle panel) and BAFF (TNFSF13B, right panel) are shown, in relation to cancer subtypes, identified by the Pam50 geneset. **D.** A heatmap of the annotated gene expression heatmap is presented.

Furthermore, cBioPortal^®^ for Cancer Discovery ^23–25^ data analysis from a large breast cancer study (2509 samples, MetaBRIC study) ^26,27^ show that BAFF, APRIL, and BCMA have a significant tendency to co-express with androgen receptor AR (p<0.001 for BAFF and APRIL, p=0.012 for BCMA) further confirming the association with androgen (Figure 6 C and D and Supplementary Figure 4). Significant correlations were also found for the stemness markers we have tested in the breast cancer cell lines.

## Discussion

Our understanding of breast cancer physiology has expanded in recent years. In addition to the molecular analysis and identification of different breast cancer subtypes ^28,29^, recent data reveal a biological evolution of breast cancer cells from the primary to the metastatic site of the disease, influencing the disease progression ^30^. This evolution has been recently suggested as a possible novel therapeutic target in breast cancer ^31^.

A major element of breast cancer molecular and phenotypic evolution is the differentiation of stem cells from their niches ^3,5,30^ and the cross-talk with the epithelial, stromal and immune components of the niche or surrounding microenvironment ^2,4,5,30–32^. However, another element of this heterogeneity has been recently identified and is related to the effects of administered treatment to the transformation of cancer epithelial cells ^33^, including the emergence of cell pluripotency. Recently, we have reported that short term exposure of breast cancer cells to tamoxifen may act as such an epigenetic stem-inducing stimulus ^34^. Here, we provide evidence about such an induction of pluripotency by the epithelial APRIL and BAFF, through BCMA, a mechanism which is related to extranuclear androgen effects.

The various markers currently used for breast cancer stem cells (BCSCs)^35,36^ identify slightly different subpopulations and it is, therefore, necessary to utilize more than one specific marker and/or to test for unique cancer stem cell properties such as tumor initiation (sphere formation or xenografted tumor development, etc) ^37^. Therefore, in the present work, to investigate the effect of APRIL and BAFF, we utilized multiple approaches to detect BCSCs. Our results show that both BAFF and APRIL increase the percentage of breast cancer stem cells. Indeed, when breast cancer cells were treated for 4 days with either BAFF or APRIL a two to three-fold increase in the BCSC population was observed. This finding suggests that the epigenetic changes required for the induction of this phenotype need a prolonged exposure to these agents.

In fact it was found that breast cancer cells, treated with APRIL or BAFF display a more aggressive phenotype as they exhibit: (1) Increased migration and modifications of the actin cytoskeleton; (2) increased expression of mesenchymal markers and signs of the acquisition of a mesenchymal phenotype that is linked to stem-like cells ^38,39^ and is associated with metastasis and a higher migratory capacity ^40,41^; (3) increased stem cell population, as assayed by BCSC autofluorescence and ALDH1A1 activity, and (4) increased mammosphere formation, with a significant increase in BCSC population. These results are in accordance with previously reported findings on APRIL promoting breast cancer lung metastasis ^18^.

An important finding supporting the observed BCSC enrichment after treatment with APRIL or BAFF is the significant acquisition of pluripotency markers KLF4, NANOG, c-MYC, and ALDH1A1 ^42–45^. These markers when properly modified, may act as transcription factors that promote a stem cell phenotype, maintain pluripotency and prevent differentiation. The fact that in the studied cell lines both BAFF and APRIL induced similar effects on these pluripotency markers at the mRNA and the protein level suggests that their action should be mediated through BCMA, a cognate receptor for both ligands ^12^. This was further supported by the inhibitory action of the specific anti-BCMA siRNA on their effects and the fact that the BCMA specific signaling involving JNK activation was found to be involved. Indeed, their effect on ALDH1A1 and KLF4 was significantly reduced in the presence of the JNK inhibitor, SP600125 or shRNAs against JNK1 and JNK2 (Figure 5C and D). Additionally, no NFκB activity was detected. Therefore, it seems that in breast cancer cells, APRIL-BCMA signal towards pluripotency is mediated by JNK, a signaling pathway also reported previously by our group for the effect of APRIL in hepatocellular carcinoma cell lines ^46^. The complete inefficiency of BAFF to modify NFκB, further suggests that, in our settings, BAFFR signaling, might be ineffective in this cell line. Moreover, we report that this APRIL-BCMA regulated effect relays on local androgen. By analyzing our previously reported transcriptome data on T47D and MDA cells ^16,47,48^, we confirmed that APRIL transcription was significantly increased after a 3h-incubation with testosterone while estradiol had only a minimal effect. Interestingly, a significantly higher change was induced by membrane-only acting testosterone (Testosterone-BSA) suggesting a membrane-initiated action, while the inhibition of the effect by anti-androgen suggests that this effect might be mediated by the classical androgen receptor, activated at the membrane level.

This androgen-induced APRIL production and subsequently, stemness, could possibly have a clinical importance in the case of breast cancer patients that are treated with aromatase inhibitors which can result at increased local androgen concentration. In order to provide a proof of principle of this hypothesis, we extracted APRIL data from a series of 37 letrozole-responders and 16 non-responders before and after therapy (GEO-deposited GDS3116 study) ^21,22^. Indeed, APRIL expression was significantly higher in the responder’s group (expected to have increased local androgen levels), while it was not changed in the non-responders, verifying our hypothesis on testosterone-control of APRIL in breast cancer.

We have also interrogated one large MetaBRIC breast cancer study ^26,27^. APRIL mRNA downregulation was evidenced in 7% of cases, confirming our previous report on protein expression ^17^. Moreover, it is evident that BAFF, APRIL, and BCMA tend to co-occur with AR and stemness markers. They are preferentially expressed in claudin-low tumors, that display a highly aggressive behavior.

Our study presents a number of limitations. First, only two breast cancer lines are tested. Larger studies in more breast cancer cell lines and in vivo models are required to validate our data. However, evidence from a number of clinical studies, presented above, referring to treated patients provide a degree of validation in patients’ data. Our omics approaches based on a retrospective analysis of publicly available data also supports our results. Future prospective studies could provide more refined data to the mechanism presented here. The clinical implications could be potentiated, as aromatase inhibition is widely used to treat ER-positive tumors and AR profiling is part of cutting-edge approaches in breast cancer.

## Conclusions

Our findings clearly indicate that both BAFF and APRIL have the ability to increase the percentage of breast cancer stem cells through BCMA-JNK mediation, pointing out for the first time that APRIL and BAFF not only modify breast cancer cell proliferation but they can also contribute to the re-formation of the tumor. Additionally, they provide evidence for a new possible mechanism of therapy resistance that involves increased stemness by high APRIL levels as a result of the accumulation of androgen that could occur in aromatase inhibitors treated patients. They further provide evidence that, in order to establish optimal personalized, immune-related therapies in breast cancer patients, one should, in addition to the targeting of the stroma and cancer-infiltrating immune cells ^8^, also investigate and target tumor cells, as was recently reported for another member of the TNFRSF, TNFR2 ^10,11^.

## Methods

### Cell cultures and chemicals

The T47D breast cancer cell line and MDA-MB-231 were purchased from DSMZ (Braunschweig, Germany), and were cultured in RPMI supplemented with 10% fetal bovine serum (FBS), at 37 °C, 5% CO_2_. All media were purchased from Invitrogen (Carlsbad, USA) and all chemicals from Sigma (St. Louis, MO) unless otherwise stated.

### RT-PCR

RT-PCR was performed as described previously ^46^. Positive controls were run in parallel with samples included in the study. We used adipose-derived mesenchymal stem cells as positive controls of BAFF and APRIL ^15^, isolated human lymphocytes as positive controls for BAFFR and TACI, and HepG2 cells for BCMA^46^. The primers used for the study were as follows: **BAFF** forward: 5′-TTC TAG GGC ACT TCC CCT TT-3′, reverse: 5′-CTC AAG ACT GCT TGC AAC TGA-3′; **APRIL** forward: 5′-TCT CCT TTT CCG GGA TCT CT-3′, reverse: 5′-CCA GAA TGG GGA AGG GTA TC-3′; **BAFFR** forward: 5′-AGG ACG CCC CAG AGC C-3′, reverse: 5′-AGT GTC TGT GCT TCT GCA GG-3′; **TACI** forward: 5′-AGT GAA CCT TCC ACC AGA GC-3′, reverse: 5′-CTC TTC TTG AGG AAG CAG GC-3′; **BCMA** forward: 5′-GTC AGC GTT ATT GTA ATG CAA GTG T-3′, reverse: 5′-TCT TTT CCA GGT CAA TGT TAG CC-3′; **18S** RNA forward: 5′-ATG GTC AAC CCC ACC GTG T-3′, reverse: 5′-TTC TGC TGT CTT TGG AAC TTT GTC-3′. All primers were selected from qPrimer Depot (qPrimerDepot, http://primerdepot.nci.nih.gov) and synthesized by VBC Biotech (Vienna, Austria).

### Real Time PCR

Total RNA was isolated with the Nucleospin RNA II isolation kit (Macherey-Nagel EURL, Fr). The absence of DNA was verified by PCR for G3PDH. One μg of RNA was subjected to ABI High Capacity cDNA reverse transcription kit (Applied Biosystems, Foster City, CA). Real-time PCR with SYBR Green was performed with DyNAmo SYBR Green qPCR Kit (Finnzymes, Oy, Finland), using the StepOnePlus™ System (Applied Biosystems), at 95°C for 3 min followed by 40 cycles of 95°C for 15 sec, 60°C for 30 sec, 72°C for 60 sec. PCR reactions were performed using the following primer pairs (synthesized by Eurofins Genomics): **c-MYC**: forward CACCGAGTCGTAGTCGAGGT and reverse TTTCGGGTAGTGGAAAACCA **SOX-2**: forward AGG AGC CTT CCT TTT CCA GA and reverse CGACGAGAGCTCCTACCAAC, **ALDH1A1**: forward TCCTCCTCAGTTGCAGGATT and reverse GCACGCCAGACTTACCTGTC, **KLF4**: forward GTCAGTTCATCTGAGCGGG and reverse AGAGTTCCCATCTCAAGGCA. The number of cycles used was optimized to fall within the linear range of PCR amplification. Changes were normalized according to cyclophilin A expression (forward: ATG GTC AAC CCC ACC GTG T and reverse: TTC TGC TGT CTT TGG AAC TTT GTC).

### Detection of cancer stem cells

***Autofluorescence based detection***: Cells after treatment with APRIL or BAFF (100ng/ml) for 4 days were detached by trypsin-EDTA from the culture plate and centrifuged (800g 10min). The pellet was re-suspended in PBS+2% FBS at a cell concentration of 1×10^6^ cells/ml. They were analyzed by flow cytometry (Attune^®^ Acoustic Focusing Cytometer, Applied Biosystems) at a cell population of at least 20000 at 488 (580/30)/488(530/40 (BL2-A/BL1-A). Dot Blot Diagram taking advantage the finding of Miranda-Lorenzo et al. 2014 ^20^ that cancer stem cells exhibit a higher level of autofluorescence.

***Aldehyde Dehydrogenase activity based detection***: Stem cells, that have the characteristic to express high levels of the enzyme aldehyde dehydrogenase (ALDH) were detected by the use of ALDEFLUOR™ kit (Stem Cell Technologies Inc., Vancouver, Canada). According to the manufacturer’s instructions, a cell suspension of 1×10^6^ cells/ml assay buffer, untreated or after treatment with APRIL or BAFF (100ng/ml) for 4 days, was incubated (45 min, at 37°C) with the ALDEFLUOR™ Reagent which is a fluorescent substrate for ALDH with (control sample) or without (test sample) the specific ALDH inhibitor dimethylamino benzaldehyde (DEAB). The fluorescent reaction product, that is retained within the cells and is proportional to the activity of ALDH, was measured by flow cytometry (Attune^®^ Acoustic Focusing Cytometer, Applied Biosystems). Data acquisition was performed using identical instrument settings for each test and control sample on a population of 20000 cells in a SSC-H/BL1-H Dot Blot Diagram.

### Mammosphere formation

Mammosphere formation was assayed as previously described ^49^. For primary mammosphere generation, cells (70-80% confluent) were detached by trypsinization, centrifuged at 580 x g for 2 min, resuspended in 2 ml of ice-cold PBS and passed several times through a 25 G syringe needle. Then cells were seeded in 6-well plates, coated with 1ml/well of 1.2% poly-(2-hydroxyethyl methacrylate (pHEMA) solution in absolute ethanol, for 48h at 40°C (600 cells/cm^2^). Cells were incubated in the presence of APRIL or BAFF (100ng/ml), testosterone-BSA (10^−6^M) or vehicle, in a humidified atmosphere at 37°C and 5 % CO2, for 7-12 days, without disturbing the plates or replenishing the medium. The number of mammospheres (primary generation, defined as a cellular mass of at least 10 cells more than 50μm in size) was counted, with a Leica DMIRE2 inverted microscope, at 40X magnification. For the generation of secondary mammospheres, primary mammospheres were collected, washed twice with 1ml PBS, centrifuged at 115g for 5 minutes and trypsinized with 300 μl of 0.5 % trypsin/0.2 % EDTA at 37°C for 2 min. After trypsin neutralization with 1ml of serum-containing medium and centrifugation at 580 g for 5 min, single cells from were resuspended in 200 μl of ice-cold PBS and seeded in pHEMA coated 6-well plates (600 cells/cm^2^). Cells were incubated for 7 additional days and secondary mammospheres were counted as described above.

### Immunocytochemistry

For the detection of immunoreactive proteins in breast cancer cells, treated or not (control) with APRIL or BAFF (100ng/ml) for 4 days, cells were fixed with 2% paraformaldehyde. Afterwards, they were incubated at room temperature for one hour with 3% BSA in TBS, and then overnight at 4 °C with the primary antibody (c-MYC 1/600, ALDH1A1 1/800, SOX-2 1/100 all from Santa-Cruz, and NANOG 1/100 from eBiosciences). For the detection of antibody binding, the UltraVision LP Detection System: HRP Polymer Quanto (Thermo Scientific, Cheshire, UK) was used, with 3’3-Diaminobenzidine (DAB) as chromogen. Stained cells were lightly counterstained with Harris hematoxylin for 10 s, hydrated and mounted in Permount (Fisher Scientific, Fair Lawn, NJ). Internal controls for specificity of immunostaining included replacement of primary antibody with non-specific serum (negative). Slides were evaluated for the presence and the intensity of staining (quantified using ImageJ).

### EMT transition

The EMT status of cells was determined by measuring the ratios of vimentin/keratins expression levels, using image analysis after laser scanning microscopy ^50^ and FACS analysis. Cells in cytospin preparations (used for microscopy) and in suspension (used for microscopy and FACS) were immunostained with primary antibodies for keratins 8, 18, 19 (mouse monoclonal antibodies, A45-B/B3, R002A, Micro-met AG, Munich, Germany) and vimentin (polyclonal antibody, Santa Cruz Biotechnology, sc-7558) and anti-mouse and anti-rabbit secondary antibodies labeled with Alexa 488 (Invitrogen) and CF555 (red staining, Biotium) dyes respectively. Untreated and treated cells from cytospin preparations and immunostained suspended cells attached on alcian blue coated coverslips were analyzed by confocal (Leica SP) microscopy. To prevent any signal interference (green and red) generated by the different emission spectra, the detection of each one of the markers was performed by sequential laser confocal scan. Fixed confocal settings were used for all specific measurements. Images were taken from 50 cells of each treatment and were stored electronically. To quantify the fluorescence intensity of vimentin and keratins, the images were subjected to Java-based image processing with the use of Image J program (NIH). In parallel, cells immunostained in suspension were also analyzed by FACS (Attune^®^ Acoustic Focusing Cytometer, Applied Biosystems) at a cell population of at least 20000 cells.

### Wound healing assay

Cells were seeded in12-well plates and cultured until a monolayer was formed. At that point, cells were incubated for 1 hour with mitomycin C (10 μg/ml), so as to prevent further cell proliferation. Afterwards, a thin wound was drawn in the middle of each well using a sterile micropipette tip and cells were washed with PBS in order to remove any remaining cell debris. Fresh medium containing BAFF or APRIL (100 ng/ml) with or without Testosterone-BSA (10^−6^ M), was added. The subsequent colonization of the denuded area was photographed with an inverted microscope (DM IRE2, Leica), at different time-intervals (at time 0, just before adding the tested agents 24 hours and 48 hours afterward, see Results for details) always at predefined points of the well. The photographs were analyzed and the covered distance was measured, and compared to the control cells.

### Actin cytoskeleton staining and visualization

Cells were grown on 8-well chamber slides. After incubation with the different agents for 10 min, cells were washed with PBS twice, fixed with 4% paraformaldehyde in PBS for 10 minutes at room temperature, permeabilized with 0.5% Triton X-100 for 10 min and incubated in blocking buffer (2% BSA in PBS) for 15 min. Actin cytoskeleton was visualized by rhodamine-phalloidin staining (1:400 in PBS containing 0.2% BSA) for 45 min. Specimens were analyzed in a Leica SP confocal microscope.

### Knock down experiments (BCMA, JNK)

T47D cells were plated in six-well plates (2.3 10^5^ cells/well) and left to adhere for 24 h. The medium was changed, and transfection with siRNA for BCMA (NM_001192, Sigma-Aldrich) or shRNA against JNK1 or JNK2 prepared by our group ^46^ for JNK was performed with a standard Lipofectamine 2000 protocol (Invitrogen; 0.8 mg DNA, 1 ml Lipofectamine 2000 in Optimem medium, for each well of a 24-well plate, scaling up for 6-well plates). For the later transfection efficiency was 85%, as estimated by counting GFP-positive cells.

### JNK inhibition

T47D cells were cultured in six-well plates and treated with APRIL or BAFF (100ng/ml) for 4 days in the presence of the specific JNK inhibitor, SP600125 (10μM).

### NFk-B activation assay

Cells were cultured in 24-well plate, and were transfected with 0.2μg/well of pNFκB-Luc plasmid (Clontech, Mountain View, CA), carrying NFκB response elements, in front of the 5’ end of the firefly luciferase gene, together with 0.2μg/well of a Renilla luciferase vector (pRL-CMV, Promega, Fitchburg, WI), using Lipofectamine 2000 (Invitrogen, 1 ml per well) in Optimem medium. Cells were incubated for 24h and then treated with BAFF or APRIL for 24 hours. Luciferase activity was assayed with a Dual-Luciferase Reporter 1000 Assay System (Promega, Fitchburg, WI), in a Berthold FB12 Luminometer (Bad Wildbad Germany).

### Data from public databases

Data for BAFF, APRIL, BCMA, AR, SOX2, KLF4, ALDH1A, NANOG2 were retrieved from cBioPortal^®^ for Cancer data analysis http://www.cbioportal.org/data_sets.jsp (version 1.4.2 snapshot) from the MetaBRIC breast cancer study (2509 samples) ^26,27^. Co-occurrence and/or mutual exclusivity, as well as correlation with breast cancer subtype and ER status, were interrogated. In addition, APRIL trascriptome data were extracted from the GEO-deposited study GDS3116 (https://www.ncbi.nlm.nih.gov/geo/) referring to 53 letrozole-treated patients was interrogated for the expression of APRIL, before and 3-months after therapy ^21,22^. (MetaBRIC study).

### Statistical analysis

Statistics were performed by the use of SPSS v21 (SPSS/IBM Inc, Chicago, IL) and Prism v6.05 (GraphPad Inc, La Jolla, CA). Parametric tests were used, as appropriate and a p<0.05 level was retained for significance.

## Authors’ contributions

VP, GN, AT, EC, and MK conceived and designed the experiments and wrote the paper. VP, GN, PA, KA, FK, NK, KK, EK, HP and PT performed the experiments and analyzed the data. All authors approved the manuscript.

## References

1 Batlle, E. & Clevers, H. Cancer stem cells revisited. Nat Med 23, 1124–1134, doi:10.1038/nm.4409 (2017).

2 Lloyd-Lewis, B., Harris, O. B., Watson, C. J. & Davis, F. M. Mammary Stem Cells: Premise, Properties, and Perspectives. Trends Cell Biol 27, 556–567, doi:10.1016/j.tcb.2017.04.001 (2017).

3 Van Keymeulen, A. et al. Lineage-Restricted Mammary Stem Cells Sustain the Development, Homeostasis, and Regeneration of the Estrogen Receptor Positive Lineage. Cell Rep 20, 1525–1532, doi:10.1016/j.celrep.2017.07.066 (2017).

4 Simoes, B. M., Alferez, D. G., Howell, S. J. & Clarke, R. B. The role of steroid hormones in breast cancer stem cells. Endocr Relat Cancer 22, T177–186, doi:10.1530/ERC-15-0350 (2015).

5 Sreekumar, A., Roarty, K. & Rosen, J. M. The mammary stem cell hierarchy: a looking glass into heterogeneous breast cancer landscapes. Endocr Relat Cancer 22, T161–176, doi:10.1530/ERC-15-0263 (2015).

6 Dyck, L. & Mills, K. H. G. Immune checkpoints and their inhibition in cancer and infectious diseases. Eur J Immunol 47, 765–779, doi:10.1002/eji.201646875 (2017).

7 Galluzzi, L., Buque, A., Kepp, O., Zitvogel, L. & Kroemer, G. Immunogenic cell death in cancer and infectious disease. Nat Rev Immunol 17, 97–111, doi:10.1038/nri.2016.107 (2017).

8 Dougan, M. & Dougan, S. K. Targeting Immunotherapy to the Tumor Microenvironment. J Cell Biochem, doi:10.1002/jcb.26005 (2017).

9 Anderson, K. G., Stromnes, I. M. & Greenberg, P. D. Obstacles Posed by the Tumor Microenvironment to T cell Activity: A Case for Synergistic Therapies. Cancer Cell 31, 311–325, doi:10.1016/j.ccell.2017.02.008 (2017).

10 Chen, X. & Oppenheim, J. J. Targeting TNFR2, an immune checkpoint stimulator and oncoprotein, is a promising treatment for cancer. Sci Signal 10, doi:10.1126/scisignal.aal2328 (2017).

11 Torrey, H. et al. Targeting TNFR2 with antagonistic antibodies inhibits proliferation of ovarian cancer cells and tumor-associated Tregs. Sci Signal 10, doi:10.1126/scisignal.aaf8608 (2017).

12 Aggarwal, B. B., Gupta, S. C. & Kim, J. H. Historical perspectives on tumor necrosis factor and its superfamily: 25 years later, a golden journey. Blood 119, 651–665, doi:10.1182/blood-2011-04-325225 (2012).

13 Rihacek, M. et al. B-Cell Activating Factor as a Cancer Biomarker and Its Implications in Cancer-Related Cachexia. Biomed Res Int 2015, 792187, doi:10.1155/2015/792187 (2015).

14 Moreaux, J., Veyrune, J. L., De Vos, J. & Klein, B. APRIL is overexpressed in cancer: link with tumor progression. BMC Cancer 9, 83, doi:10.1186/1471-2407-9-83 (2009).

15 Alexaki, V. I. et al. Adipocytes as immune cells: differential expression of TWEAK, BAFF, and APRIL and their receptors (Fn14, BAFF-R, TACI, and BCMA) at different stages of normal and pathological adipose tissue development. J Immunol 183, 5948–5956, doi:10.4049/jimmunol.0901186 (2009).

16 Pelekanou, V. et al. The estrogen receptor alpha-derived peptide ERalpha17p (P(295)-T(311)) exerts pro-apoptotic actions in breast Cancer cells in vitro and in vivo, independently from their ERalpha status. Mol Oncol 5, 36–47, doi:10.1016/j.molonc.2010.11.001 (2011).

17 Pelekanou, V. et al. Expression of TNF-superfamily members BAFF and APRIL in breast cancer: immunohistochemical study in 52 invasive ductal breast carcinomas. BMC Cancer 8, 76, doi:10.1186/1471-2407-8-76 (2008).

18 Garcia-Castro, A. et al. APRIL promotes breast tumor growth and metastasis and is associated with aggressive basal breast cancer. Carcinogenesis 36, 574–584, doi:10.1093/carcin/bgv020 (2015).

19 Brooks, M. D. & Wicha, M. S. Tumor twitter: cellular communication in the breast cancer stem cell niche. Cancer Discov 5, 469–471, doi:10.1158/2159-8290.CD-15-0327 (2015).

20 Miranda-Lorenzo, I. et al. Intracellular autofluorescence: a biomarker for epithelial cancer stem cells. Nat Methods 11, 1161–1169, doi:10.1038/nmeth.3112 (2014).

21 Miller, W. R. et al. Changes in breast cancer transcriptional profiles after treatment with the aromatase inhibitor, letrozole. Pharmacogenet Genomics 17, 813–826, doi:10.1097/FPC.0b013e32820b853a (2007).

22 Miller, W. R. & Larionov, A. Changes in expression of oestrogen regulated and proliferation genes with neoadjuvant treatment highlight heterogeneity of clinical resistance to the aromatase inhibitor, letrozole. Breast Cancer Res 12, R52, doi:10.1186/bcr2611 (2010).

23 Cerami, E. et al. The cBio cancer genomics portal: an open platform for exploring multidimensional cancer genomics data. Cancer Discov 2, 401–404, doi:10.1158/2159-8290.CD-12-0095 (2012).

24 Gao, J. et al. Integrative analysis of complex cancer genomics and clinical profiles using the cBioPortal. Sci Signal 6, pl1, doi:10.1126/scisignal.2004088 (2013).

25 Cheng, G. Z. et al. Twist transcriptionally up-regulates AKT2 in breast cancer cells leading to increased migration, invasion, and resistance to paclitaxel. Cancer Res 67, 1979–1987, doi:10.1158/0008-5472.CAN-06-1479 (2007).

26 Pereira, B. et al. The somatic mutation profiles of 2,433 breast cancers refines their genomic and transcriptomic landscapes. Nat Commun 7, 11479, doi:10.1038/ncomms11479 (2016).

27 Curtis, C. et al. The genomic and transcriptomic architecture of 2,000 breast tumours reveals novel subgroups. Nature 486, 346–352, doi:10.1038/nature10983 (2012).

28 Sorlie, T. et al. Gene expression patterns of breast carcinomas distinguish tumor subclasses with clinical implications. Proc Natl Acad Sci U S A 98, 10869–10874, doi:10.1073/pnas.191367098 (2001).

29 Paquet, E. R. & Hallett, M. T. Absolute assignment of breast cancer intrinsic molecular subtype. J Natl Cancer Inst 107, 357, doi:10.1093/jnci/dju357 (2015).

30 Yates, L. R. et al. Genomic Evolution of Breast Cancer Metastasis and Relapse. Cancer Cell 32, 169–184 e167, doi:10.1016/j.ccell.2017.07.005 (2017).

31 Amirouchene-Angelozzi, N., Swanton, C. & Bardelli, A. Tumor Evolution as a Therapeutic Target. Cancer Discov, doi:10.1158/2159-8290.CD-17-0343 (2017).

32 Wang, X. et al. Breast tumors educate the proteome of stromal tissue in an individualized but coordinated manner. Sci Signal 10, doi:10.1126/scisignal.aam8065 (2017).

33 Giltnane, J. M. et al. Genomic profiling of ER+ breast cancers after short-term estrogen suppression reveals alterations associated with endocrine resistance. Sci Transl Med 9, doi:10.1126/scitranslmed.aai7993 (2017).

34 Notas, G. et al. Tamoxifen induces a pluripotency signature in breast Cancer Cells and human tumors. Mol Oncol 9, 1744–1759, doi:10.1016/j.molonc.2015.05.008 (2015).

35 Al-Hajj, M., Wicha, M. S., Benito-Hernandez, A., Morrison, S. J. & Clarke, M. F. Prospective identification of tumorigenic breast cancer cells. Proc Natl Acad Sci U S A 100, 3983–3988, doi:10.1073/pnas.0530291100 (2003).

36 Iqbal, J., Chong, P. Y. & Tan, P. H. Breast cancer stem cells: an update. J Clin Pathol 66, 485–490, doi:10.1136/jclinpath-2012-201304 (2013).

37 Kok, M. et al. Mammosphere-derived gene set predicts outcome in patients with ER-positive breast cancer. J Pathol 218, 316–326, doi:10.1002/path.2544 (2009).

38 Santisteban, M. et al. Immune-induced epithelial to mesenchymal transition in vivo generates breast cancer stem cells. Cancer Res 69, 2887–2895, doi:10.1158/0008-5472.CAN-08-3343 (2009).

39 Mani, S. A. et al. The epithelial-mesenchymal transition generates cells with properties of stem cells. Cell 133, 704–715, doi:10.1016/j.cell.2008.03.027 (2008).

40 Tsai, J. H. & Yang, J. Epithelial-mesenchymal plasticity in carcinoma metastasis. Genes Dev 27, 2192–2206, doi:10.1101/gad.225334.113 (2013).

41 Tiwari, N., Gheldof, A., Tatari, M. & Christofori, G. EMT as the ultimate survival mechanism of cancer cells. Semin Cancer Biol 22, 194–207, doi:10.1016/j.semcancer.2012.02.013 (2012).

42 Chambers, I. & Tomlinson, S. R. The transcriptional foundation of pluripotency. Development 136, 2311–2322, doi:10.1242/dev.024398 (2009).

43 Rodriguez-Pinilla, S. M. et al. Sox2: a possible driver of the basal-like phenotype in sporadic breast cancer. Mod Pathol 20, 474–481, doi:10.1038/modpathol.3800760 (2007).

44 Karoubi, G., Gugger, M., Schmid, R. & Dutly, A. OCT4 expression in human non-small cell lung cancer: implications for therapeutic intervention. Interact Cardiovasc Thorac Surg 8, 393–397, doi:10.1510/icvts.2008.193995 (2009).

45 Jeter, C. R. et al. Functional evidence that the self-renewal gene NANOG regulates human tumor development. Stem Cells 27, 993–1005, doi:10.1002/stem.29 (2009).

46 Notas, G. et al. APRIL binding to BCMA activates a JNK2-FOXO3-GADD45 pathway and induces a G2/M cell growth arrest in liver cells. J Immunol 189, 4748–4758, doi:10.4049/jimmunol.1102891 (2012).

47 Notas, G. et al. Whole transcriptome analysis of the ERalpha synthetic fragment P295-T311 (ERalpha17p) identifies specific ERalpha-isoform (ERalpha, ERalpha36)-dependent and - independent actions in breast cancer cells. Mol Oncol 7, 595–610, doi:10.1016/j.molonc.2013.02.012 (2013).

48 Kampa, M. et al. Early membrane initiated transcriptional effects of estrogens in breast cancer cells: First pharmacological evidence for a novel membrane estrogen receptor element (ERx). Steroids 77, 959–967, doi:10.1016/j.steroids.2012.02.011 (2012).

49 Shaw, F. L. et al. A detailed mammosphere assay protocol for the quantification of breast stem cell activity. J Mammary Gland Biol Neoplasia 17, 111–117, doi:10.1007/s10911-012-9255-3 (2012).

50 Polioudaki, H. et al. Variable expression levels of keratin and vimentin reveal differential EMT status of circulating tumor cells and correlation with clinical characteristics and outcome of patients with metastatic breast cancer. BMC Cancer 15, 399, doi:10.1186/s12885-015-1386-7 (2015).

